# From spreadsheet lab data templates to knowledge graphs: A FAIR data journey in the domain of AMR research

**DOI:** 10.1101/2024.07.18.604030

**Authors:** Yojana Gadiya, Tooba Abbassi-Daloii, Vassilios Ioannidis, Nick Juty, Claus Stie Kallesøe, Marie Attwood, Manfred Kohler, Philip Gribbon, Gesa Witt

**Author notes:** **Corresponding author:** Yojana Gadiya, Fraunhofer Institute for Translational Medicine and Pharmacology. These authors contributed equally to the work (Yojana Gadiya, Tooba Abbassi-Daloii, Gesa Witt).

## Abstract

While awareness of FAIR (Findable, Accessible, Interoperable, and Reusable) principles has expanded across diverse domains, there remains a notable absence of impactful narratives regarding the practical application of FAIR data. This gap is particularly evident in the context of *in-vitro* and *in-vivo* experimental studies associated with the drug discovery and development process. Despite the structured nature of these data, reliance on classic methods such as spreadsheet-based visualization and analysis has limited the long-term reuse opportunities for such datasets. In response to this challenge, our work presents a representative journey towards FAIR data, characterized by structured, conventional spreadsheet-based lab data templates and the adoption of a knowledge graph framework for breaking data silos in the field of early antimicrobial resistance research. Here, we illustrate a tailored application of a “FAIRification framework” facilitating the practical implementation of FAIR principles. By showcasing the feasibility and benefits of transitioning to FAIR data practices, our work aims to encourage broader adoption and integration of FAIR principles within a research lab setting.

## Introduction

The FAIR (Findable, Accessible, Interoperable, and Reusable) principles offer essential guidance to enhance the inherent value and reusability of data ^1^. They specify a set of characteristics that data should possess, encompassing not only the data itself but also the resources that contribute to its creation, such as algorithms, workflows, and the infrastructures hosting the data. Moreover, by adhering to the FAIR principles, researchers and organizations can significantly enhance the efficiency of data sharing and facilitate collaboration, advance scientific discovery, enhance visibility and recognition, promote transparency and reproducibility, and maximize the value of investments in data generation and curation. This underscores the importance of FAIR data in research, which has been widely promoted across different domains. However, the extent of its awareness and implementation varies. For instance, a study commissioned by the European Research Data Landscape revealed that while approximately 30% of participating researchers had heard about the FAIR principles, many were not fully familiar with their implications ^2^. Additionally, nearly 40% of respondents had never encountered these principles before. This has prompted a call for action and attention across various fields to identify the reasons for this discrepancy and to cultivate a more informed community around FAIR principles.

One of the key factors that has contributed to the difficulty in understanding FAIR principles is its implementation ^3^. While most researchers grasp the foundational concepts and significance of FAIR data, the absence of detailed technical guidance poses a significant barrier. Researchers require relevant case studies showing the benefits of FAIR in their respective research domains, as well as hands-on guidance and systematic (non-disruptive) processes to effectively translate the aspirational FAIR principles into actionable steps. FAIR principles emphasize the importance of making data findable through publication in established repositories, ensuring accessibility via open-access licenses, promoting interoperability through the adoption of standardized formats, and enabling reusability by facilitating the seamless integration of diverse resources. However, the actual implementation of these principles can vary depending on the specific needs, goals, and constraints of different research communities, disciplines, and projects ^4-6^. What works well in one domain may not be directly applicable or suitable for another. Therefore, flexibility and adaptation are key when applying FAIR principles to diverse datasets across disparate research contexts. The limited support and lack of monitoring of the adoption and implementation of FAIR principles are additional barriers to their acceptance. The implementation of these principles can be time-consuming and complex, often requiring collaborations among multiple stakeholders and domain experts to develop a detailed implementation plan ^7-10^.

To address these challenges and promote FAIR data practices, various initiatives have employed different but complementary strategies, for instance, RDMKit (https://rdmkit.elixir-europe.org/) cultivates FAIR data management on a global scale, FAIRSharing.org (https://fairsharing.org/) identify FAIR data champions, and Pistoia Alliance (https://www.pistoiaalliance.org/) encourage adoption across an industrial context. Each contributes in their own way to promote, educate, and assist researchers in understanding FAIR data management. Alongside these sustainable communities, there is a surging growth of grant and funding applications from governing bodies like the Federal Ministry of Education and Research Germany (BMBF), Innovative Health Initiative (IHI) (formerly Innovative Medicines Initiative (IMI)), European Open Science Cloud (EOSC) and European Commission (EC) among others, seeking to promote domain adoption and implementation of FAIR services ^11-14^. Currently, the adoption of FAIR principles is becoming increasingly mandated by most, if not all, funding agencies.

Of all the life science domains, the impact of FAIR data in the context of drug discovery and development cannot be ignored ^15-21^. The drug discovery process involves the accumulation of vast amounts of experimental and computational data, making it akin to a data warehouse of information. Adhering to FAIR principles can harness and repurpose this wealth of data, mitigating the need for redundant experiments and conserving valuable time and resources. Therefore, embracing FAIR principles within this domain is not only crucial but also indispensable for optimizing research outcomes and expediting the development of innovative therapeutics. Frontiers in this direction include initiatives at Roche and AstraZeneca, which have spearheaded an internal FAIRification movement ^22,23^. Their objective is to standardize legacy clinical data to the Clinical Data Interchange Standards Consortium (CDISC) format and to catalog this FAIR data for internal use. In doing so, they have successfully developed machine learning-based prediction models for advancing drug discovery ^24^. Likewise, the Federation of Imaging Data for Life Sciences (FIDL), a public-private collaboration project, involves the FAIRification of biomedical imaging data by definition of standards and ontologies to represent data and metadata ^25^.

Here, we examine one of the life science domains which urgently requires effective FAIR data management. Antimicrobial resistance (AMR) is a pressing global health concern characterized by microorganisms evolving mechanisms to withstand the effects of antimicrobial drugs, rendering them ineffective in treating infections. This situation poses a profound threat to public health. To combat this, the IMI has launched multiple projects that target generic antimicrobial drug discovery, such as COMBACTE-NET (https://www.combacte.com/about/about-combacte-net-detail/) and ND4BB (https://www.imi.europa.eu/projects-results/project-factsheets/nd4bb), as well as drug discovery specific projects, such as GNA NOW (https://amr-accelerator.eu/project/gna-now/) with focus on Gram-negative strains or TRIC-TB (https://amr-accelerator.eu/project/tric-tb/) for *Mycobacterium tuberculosis* resistant strain. Addressing AMR requires a multifaceted approach involving improved surveillance, responsible antibiotic stewardship, development of new antimicrobial agents, and public education initiatives to raise awareness of the importance of prudent antibiotic use. Among these approaches, the process of developing new drugs is notably complex and time-consuming, underscoring the critical role of retrospectively generated data in enhancing the likelihood of clinical success. Hence, there is a growing realization that adopting FAIR data formats for research data storage could streamline the time-to-market for new treatments. Established data standards such as the Observational Medical Outcomes Partnership Common Data Model (OMOP-CDM) and the Fast Healthcare Interoperability Resources (FHIR) are becoming increasingly employed as part of clinical data analysis workflows (https://www.ehden.eu/). However, earlier discovery stage studies still rely heavily on data producers’ specific formats. Despite databases like ChEMBL facilitating the deposition of structured data in standard formats, awareness of these resources and their underlying formats remains limited ^26^. Consequently, improving accessibility and standardization of experimental documentation in the early stages of drug discovery could significantly enhance collaboration, streamline processes, and ultimately expedite the development of urgently needed antimicrobial treatments.

In this manuscript, we focused on the FAIRification tools and implementation guidelines derived from the IMI project, FAIRplus (https://fairplus-project.eu/). The FAIRplus project, which lasted three years (2019-2022), was initiated to enhance data management and sharing practices in life sciences research. The project prioritized the refinement and adoption of FAIR data management protocols and standards alongside the creation of tools and resources designed to streamline the FAIRification process. During its lifetime, the project developed multiple FAIR data management resources, including the FAIR cookbook (https://faircookbook.elixir-europe.org/content/home.html), a living encyclopedia of ‘recipes’ to make data FAIR in the Life Sciences ^27^; the FAIR DataSet Maturity (FAIR-DSM) model (https://fairplus.github.io/Data-Maturity/), a comprehensive framework developed to assess the maturity levels of datasets, which can be used to evaluate FAIR levels before and after FAIRification efforts ^28^, and the FAIRification framework (https://w3id.org/faircookbook/FCB079), which provides practical guidance for improving the FAIRness of existing and proposed datasets ^29^. Here, we illustrate a FAIR data journey aimed at standardizing both *in-vitro* (often static with a single time point data, for e.g. minimum inhibitory concentration *MIC*) and *in-vivo* (often dynamic over multiple time points) experimental data results in a FAIR-compliant manner. While tailored to the needs of GNA NOW, the journey is designed to be adaptable across other projects with minimal effort. Through this process, we optimized a standardized lab data template for collecting data generated within various experimental labs, ensuring data harmonization with terminologies and ontologies. Additionally, we devised a workflow for data representation in the form of a knowledge graph, facilitating easy retrieval and querying of data distributed across various resources. This comprehensive approach not only enhances data accessibility and interoperability but also lays a foundation for efficient data management practices for the future.

## Results

### Tools and resources in the GNA NOW ecosystem

Similar to other collaborative IHI/IMI projects, the GNA NOW project has implemented an ecosystem of tools and resources designed to ensure efficient and secure data management, one aspect of which is to collect experimental data from various partners in a standardized manner. This ecosystem includes a cloud-based electronic lab notebook (ELN) called “Biovia Notebook Cloud” (https://www.3ds.com/products/biovia/notebook) dedicated to recording experimental results and protocols as part of the daily lab routine. In parallel, the project utilizes a scientific data repository known as “grit”, developed by the software company grit42 (https://grit42.com/), enabling storage, retrieval, querying, and comprehension of structure-activity relationships of the relevant modalities. To rationalize the integration of data into the grit platform, we developed a master “Lab Data Template” designed to collate findings from individual experiments conducted across the consortium (**Figure 1**). Practically, we created two distinct lab data templates: one for *in-vitro* studies and another for *in-vivo* studies in mice (see the **Methods section** for comprehensive details on the templates). Additionally, a close alignment between the terminologies used in the Lab Data Template and those displayed in grit was done to facilitate easy interpretation and collection of data. This process also necessitated agreement upon endpoints across the disciplines for terms such as “Colony-

**Figure 1:**
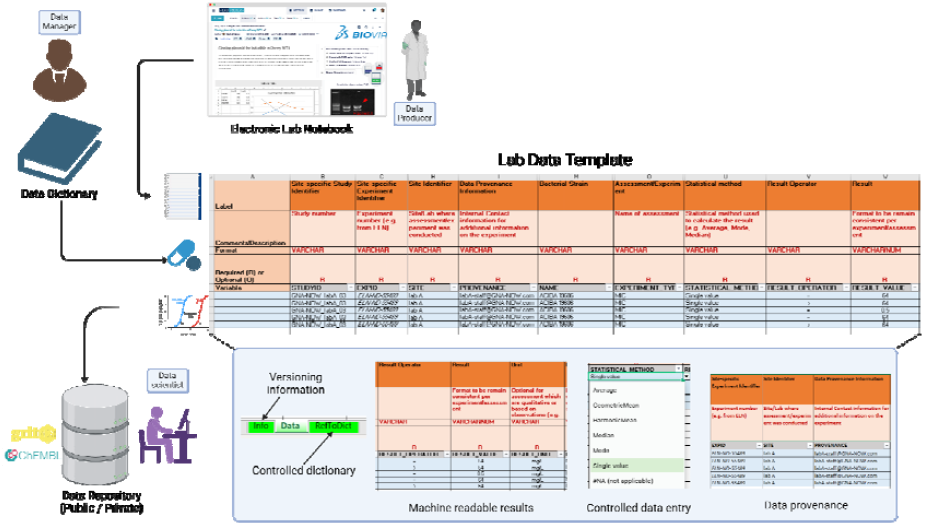
Overview of the lab data template and its correlation with the other GNA NOW tools and resources.

Forming Units” (CFU). Collectively, these template resources structure and disseminate data produced in a single laboratory to other project collaborators, including technicians and data scientists.

More broadly, establishing distinct resources to capture all types of project-generated data enhances its value but poses a significant challenge in the long term. While these modular resources are essential for capturing crucial information, they inadvertently create data and information silos. To address this issue, it is imperative to establish interconnecting links between these independent resources, enabling smooth integration and efficient browsing of data across the project’s ecosystem. This approach itself may encounter obstacles in drug discovery, particularly when collaborating with private pharmaceutical partners, due to potential restrictions on public data dissemination. This scenario was evident in the GNA NOW project, where the generated data was available only internally. It is crucial to highlight that FAIR does not necessarily mean open; rather, it allows for controlled access to data, a situation that was prominent in our case. Furthermore, the absence of a standardized data model to encompass both *in-vitro* and *in-vivo* experimental data compounded these challenges. Hence, this highlighted the need for standardized Lab Data Templates when capturing experimental data.

### Adopting the FAIRification framework for FAIR implementation

While numerous FAIR implementation frameworks are available (such as FAIR Data Point ^30^, FAIR Implementation Framework (https://fair-impact.eu/fair-implementation-framework), EOSC interoperability framework ^31^, etc.), these address predefined implementations, while the FAIRification framework is more generally applicable, with no predefined outcome ^29^. Developed as part of the FAIRplus project, this framework offers practical guidance on applying the FAIR principles to any dataset. It is user-friendly, adaptable across various domains, and supports a multi-level approach, making it an ideal approach. This framework’s four phases are: goal definition, project examination; iterative cyclical FAIRification; and post-FAIRification review (**Figure 2**). In the following sections, we will delve deeper into each phase, with a focus on the decision-making process applied to the ‘GNA NOW’ project.

**Figure 2:**
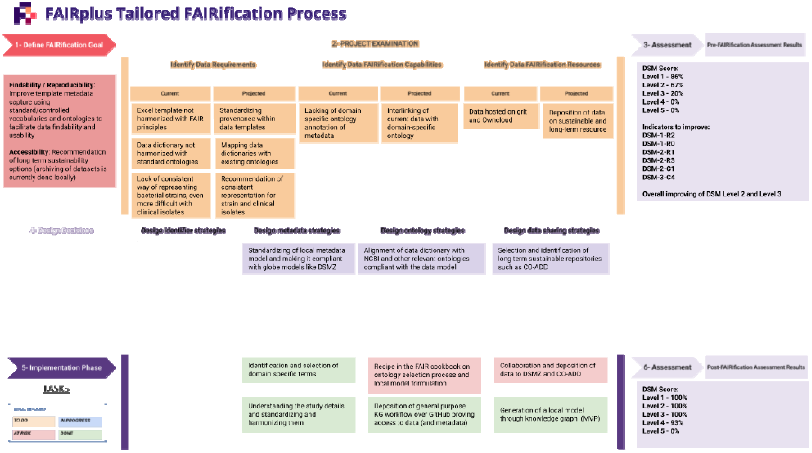
Fair assessment cycle tailored to GNA NOW using the FAIRification framework. The template was filled from Steps 1-6 based on discussions with stakeholders involving data scientists, data managers, and project leads and managers.

In goal definition (**Figure 2, Step 1**), the anticipated outcomes of FAIRification are established by identifying the intended usability of the data post-FAIRification compared to its pre-FAIRification state. Outcomes are analyzed in terms of findability (F), accessibility (A), interoperability (I), and reusability (R) of the data. However, the focus on individual aspects may vary based on the context, and not all aspects of FAIR need to be focused upon. In GNA-NOW, our overarching goal for this FAIRification effort involved inspecting the FAIRness level of the existing Lab Data Templates and identifying strategies to elevate its FAIRness. In tandem with this, like for any project, planning for sustainability solutions was key. We also wanted to explore recommendations for sustainable solutions that could aid in data dissemination for community reuse (**Figure 2, Step 1**).

Next, in the project examination phase, tasks are divided into three broad categories (**Figure 2, Step 2**). Firstly, defining the requirements for data characterization involves establishing standards and formats for the data, as well as metadata specifications. Secondly, identifying data capabilities includes documenting data access, sharing, and licensing policies, and thirdly, identifying resources such as data infrastructures for hosting and storage. The main objective here is to assess the existing strengths and limitations within these categories and anticipate how these limitations can be tackled to enhance future capabilities. It is crucial to maintain a clear focus on the primary goal while evaluating both the current and projected impacts of these categories. In GNA-NOW, we observed discrepancies between terms used in the Lab Data Templates and existing ontologies and terminologies. Furthermore, we recognized that the metadata captured through the Lab Data Template could be enhanced to encompass more detailed experimental-level information (**Figure 2, Step 2**). These improvements could eventually hold the potential to enhance experimental reproducibility, addressing a pressing issue in the field.

Following this is the iterative FAIRification cycle comprising three steps: assess, design, and implement (**Figure 2, Steps 3-5**). The initial step involves assessing the current level of FAIRness of the data, thus establishing a baseline for future comparisons. Next, the design phase is employed to pinpoint and formalize actionable tasks derived from the project examination phase in alignment with FAIRification goals. Subsequently, these tasks are executed during the implementation step. It is essential to generate a comprehensive list of tasks, recognizing that not all may be completed within a single FAIRification cycle. Note that a FAIRification cycle, similar to an agile methodology (https://agilemanifesto.org/), can be time-boxed to align with work schedules. Multiple cycles can be planned for individual projects, thereby allowing prioritized tasks to be executed first, while ‘nice to have’ implementations can be included if time allows in a cycle. This approach ensures that the FAIR journey begins with continuous improvement in mind. In the concluding phase, the post-FAIRification review (**Figure 2, Step 6**), all achievements in FAIRification undergo review to evaluate the overall success of the process. Furthermore, a final evaluation is conducted to confirm the improvement in data FAIRness.

The pre- and post-FAIRification of the dataset requires a tool to assess its FAIRness level at each stage and to iteratively revisit the assessment to evaluate improvements. Several FAIR assessment tools are available, including those listed on FAIRassist (https://fairassist.org/) ^32^. We selected the FAIR-DSM model, developed in the FAIRplus project, for this purpose ^28^. Built upon the FAIR principles, this model defines and categorizes requirements to progressively enhance the FAIRness of a given research dataset across maturity levels, ranging from level 0 to level 5, and spans three key dimensions:

a. Representation and Format - Involves questions on the data format such as JSON, Excel, PDF
b. Content and Context - Focusing on data completeness, use of ontologies, and clarity of data representation
c. Hosting environment capabilities - Covers data accessibility, such as open or closed access and programmatic access options.

At Level 0, datasets are intended for single-use purposes, whereas Level 1 represents data object-level maturity. Moving up, Level 2 pertains to project-level maturity, while Level 3 signifies community-level maturity. Level 4 extends to cross-community collaboration, and Level 5 represents enterprise-level datasets with robust data governance practices (https://fairplus.github.io/Data-Maturity/docs/Levels). Level 3 can be pragmatically defined as a “practicable” target maturity level for academic or public-private collaboration projects. At this stage, datasets are harmonized in accordance with the FAIR principles, enabling their deposition in domain-specific repositories to ensure long-term sustainability. Hence, we set our FAIRification goal to be at level 3 of maturity. The preference for the FAIR-DSM over other tools offers numerous advantages. Unlike other models, the FAIR-DSM not only assigns a score or maturity level to the dataset but also provides comprehensive explanations for why the dataset falls into a specific category or level. This transparency allows us to identify any weaknesses in the dataset and focus our efforts on addressing those areas to enhance the overall maturity level. Furthermore, its intuitive questionnaire-based format, supplemented with detailed explanations and examples in a glossary, makes it accessible even to non-FAIR experts.

Using the FAIR-DSM, we conducted a pre-FAIRification assessment to determine the current maturity level of the Lab Data Templates. **Table 1** reveals the maturity level assessment of the Lab Data Templates across the three dimensions (i.e. representation and format, content and context, and hosting environment). Whereas we found 100 % fulfillment of the FAIR-DSM criteria on level 1 in the dimensions “content and context” as well as “hosting environment”, we identified gaps in the dimension representation and format. Consequently, the assessed Lab Data Template corresponded to maturity Level 0, which means that the template had no use beyond a single purpose, which is to be used for GNA NOW internally only. The report precisely unveiled where the Lab Data Template was lacking adherence to the FAIR principles (see Zenodo dump ^39^). Notably, the absence of a standardized structure for all metadata restricted the Lab Data Template from attaining Level 1. In this context, the metadata for our Lab Data Templates refers to experiment-level requirements, such as the cell culture protocol or media conditions. In contrast, the presence of a structured format and machine-readable file, exemplified by a spreadsheet featuring unique identifiers for each experiment (leveraging the ELN identifier), proved advantageous for bolstering the content and context dimensions of the Lab Data Template. Furthermore, the hosting environment (i.e. grit platform) facilitated data retrieval and leveraged a standardized communication protocol (https://fairplus.github.io/Data-Maturity/docs/Indicators/#dsm-1-h3) with persistent identifiers, thereby elevating the maturity level within the hosting environment dimension.

**Table 1:** Pre-assessment maturity level of Lab Data Templates. The overall percentage is calculated through compliance with indicators in each category, at each level, as a proportion of the total set of indicators that are available.

| Maturity Level | Representation and Format | Content and Context | Hosting Environment | Overall |
| --- | --- | --- | --- | --- |
| Level 1 | 67 % | 100 % | 100 % | 86 % |
| **Level 2** | 50 % | 78 % | 75 % | 67 % |
| **Level 3** | 13 % | 0 % | 60 % | 20 % |
| **Level 4** | 0 % | 0 % | 0 % | 0 % |
| **Level 5** | 0 % | 0 % | 0 % | 0 % |

### Implementation strategy using knowledge graphs and ontologies

As highlighted in the preceding section, the FAIR-DSM serves as a valuable guide for the identification of the next set of tasks aimed at improving the FAIRness of the existing dataset. While the Lab Data Templates ensured structured data, the absence of ontology-linked terms and standard metadata schema hindered harmonization efforts. Thus, we identified a list of actionable tasks to aid us in achieving our FAIR goals (**Figure 2, Step 4**) whilst also setting objectives aimed at ensuring a long-term sustainable solution. One critical aspect of our sustainability approach involved identifying two bacterial-specific repositories to disseminate the GNA NOW data and thus advance community research. Firstly, the Community for Open Antimicrobial Drug Discovery (CO-ADD, https://www.co-add.org/), a crowd-sourced community data resource dedicated to screening antimicrobial compounds and libraries ^33^, and which includes bioactivity data on compounds exposed to bacterial and fungal strains. Secondly, the DSMZ’s Bacterial Diversity metadatabase (BacDive, https://bacdive.dsmz.de/), an extensive repository encompassing bacterial strain-specific genomic, proteomic, and taxonomic data, among others ^34^. BacDive serves as a centralized hub for researchers to access and explore a wealth of bacterial diversity information, facilitating deeper insights into microbial communities and their interactions. By leveraging these repositories, we aim to not only enhance data dissemination but also foster collaborative research efforts and accelerate progress in the field of antimicrobial research and beyond.

Shifting the focus to Lab Data Templates, we divided the FAIRification goal into two sub-cycles: ontology term identification and mapping in the data dictionary and independent evaluation. The former task involved standardization by identifying and mapping existing terms in the Lab Data Template to relevant ontologies and controlled vocabularies. To achieve this, we developed a data dictionary that harmonized Lab Data Template terms with domain-accepted terminology. We utilized major services such as the EMBL-EBI’s Ontology Lookup Service (https://www.ebi.ac.uk/ols4) and NCBI’s BioPortal (https://bioportal.bioontology.org/), both of which offer a comprehensive catalog of ontologies and vocabularies across domains. These tools provided the necessary resources to ensure that our terminology was consistent and aligned with widely accepted standards, thereby facilitating interoperability and enhancing the overall FAIRness of our dataset. The *in-vitro* Lab Data Template was a single-sheet Excel file with 25 columns capturing information on the study, bacterial strain, compound, and resultant activity. Among these, we identified seven categories (i.e. bacterial strain names, biomaterial, experiment type, medium, result unit, species name, and statistical method) that could benefit from linking to existing ontology terms. Analogously, the *in-vivo* Lab Data Template was distributed across four Excel sheets with 57 columns detailing information on the animal, its housing conditions, the pre- and post-treatment provided to the animal, infection strain, compound dosage, and resultant activity. In this case, we identified ten categories (i.e. animal sex, animal strain, route of administration for pre- and post-treatment, experiment type, result unit, species name, and statistical method) that could be improved through ontology term integration. Notably, due to overlaps between the columns in the two Lab Data Templates, a total of eleven categories were identified for enhancement. Since the data was generated from clinical isolates of bacterial strains, a local ontology mapping of bacterial strains was performed based on internal cataloging standards. For the remaining categories, a dedicated manual curation effort was undertaken to identify twelve ontologies that covered various aspects of the experimental data (refer to the **Methods section** for details). Interestingly, during the harmonization process, we encountered several terms that were specific to the project’s internal experiments, especially for experiment types such as inner membrane permeabilization and endpoint results like colony-forming units in lung homogenates. Consequently, a data dictionary with a hybrid setup of local (represented by the GNA NOW namespace) and global (represented by well-known namespaces like NCBI and BFO) was created. This approach ensured a comprehensive and context-appropriate data dictionary.

Next, during the evaluation of the Lab Data Template as a whole, we assessed its adherence to the tidy data principles as outlined by Wickham ^35^. According to these principles, each variable must have its own column, each observation must have its own row, and each type of observational unit must have its own table. For example, the ‘result value’ and ‘result unit’ should be in separate columns, as they are distinct variables with their own data types. By adhering to the tidy data principles, we aimed to create a robust Lab Data Template that not only meets current data management standards but also enhances the efficiency and effectiveness of data-driven research and innovation. In this context, the Lab Data Template had already aligned with these principles in certain aspects, such as recording results. The Lab Data Template captures results across multiple columns: the result operator (e.g., > or =), the result value, and the result unit. However, we found that the experimental metadata did not fully comply with the tidy data principles. To address this, we included additional columns to capture relevant experimental metadata. For instance, we split the ‘medium’ column into ‘medium pH’ and ‘medium additives’ to improve data granularity and aid reproducibility. Furthermore, we incorporated the characterization of clinically isolated bacterial strains as part of the Lab Data Template, including columns to specify the ‘source’ of samples (e.g., blood or urine) and their ‘category’ (e.g., multi-drug resistant). By making these adjustments, we ensured that the Lab Data Template not only adhered to tidy data principles but also facilitated reproducible scientific research principles.

After standardizing the Lab Data Template in multiple dimensions (at the field level with ontology linking and at the data organization level by adherence to tidy data principles), we established a structured data model to connect data across different silos present in GNA NOW. Guided by insights from the FAIR-DSM, we recognized the necessity for an infrastructure like a linked data store, specifically a knowledge graph (KG), to facilitate this transition. The initial step in setting up this KG involved defining an underlying schema to coherently connect different Lab Data Template terms and acknowledge the presence of data across multiple resources. We combined the data from both *in-vitro* and *in-vivo* Lab Data Templates to achieve comprehensive coverage of all experiments conducted within the GNA NOW consortium, as well as to track compounds in the screening workflows. By integrating information from both Lab Data Templates, we identified thirteen nodes or entities in our graph. These included specimen and bacterial strains from the *in-vivo* Lab Data Template, as well as animal species, animal groups, and study information from the *in-vitro* Lab Data Template (**Figure 3**). Overlapping data from both Lab Data Templates encompassed compound, batch, experiment type, and results. Moreover, a key focus was placed on capturing metadata for each node in the graph to facilitate the interlinking of various resources utilized in the GNA NOW project, where experiment-related data was stored. These resources comprised links to the ELN entry, housing experimental protocols for the experiment, and the grit entry, housing compound information, assay results, and their related structure-activity relationships. Despite their independence, the KG served as an overarching platform connecting information across both resources, supporting the navigation of distributed data (**Figure 4**).

**Figure 3:**
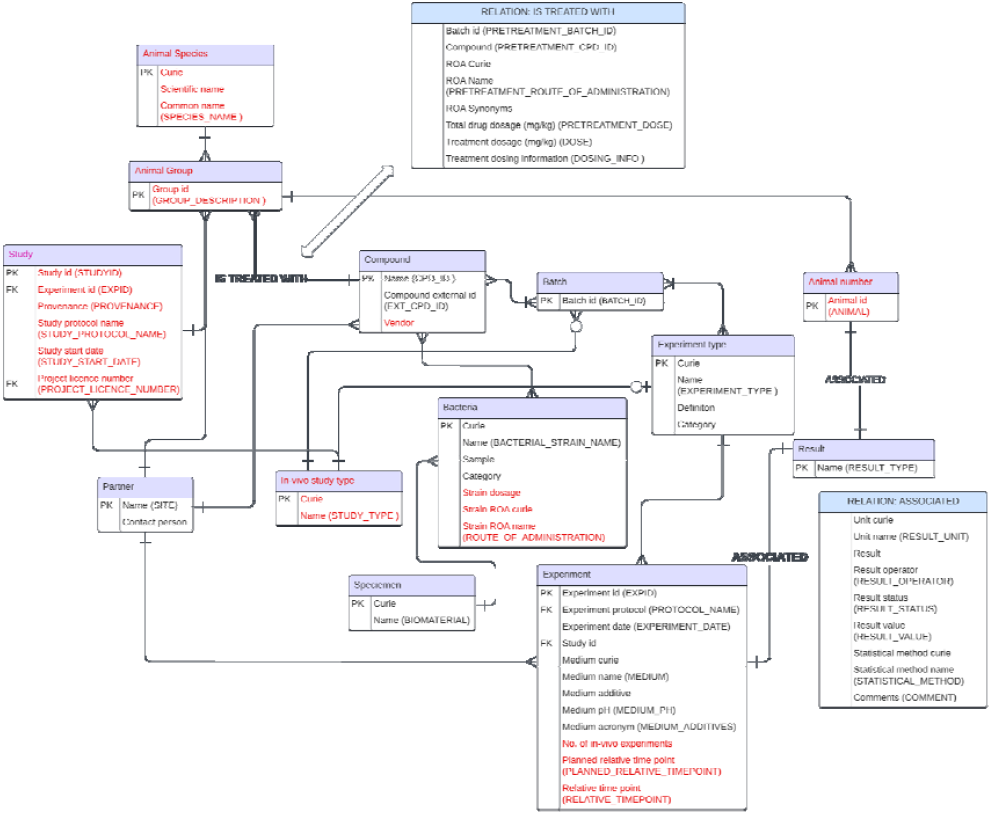
Graph database schema connecting *in-vivo* and *in-vitro* datasets. The nodes (along with their properties) in the graph are represented by purple boxes, while the relations that have metadata are represented by blue boxes. Furthermore, node (or node properties) text in black denotes data captured from the *in-vitro* Lab Data Template, whereas that in red captures data from the *in-vivo* Lab Data Template.

**Figure 4:**
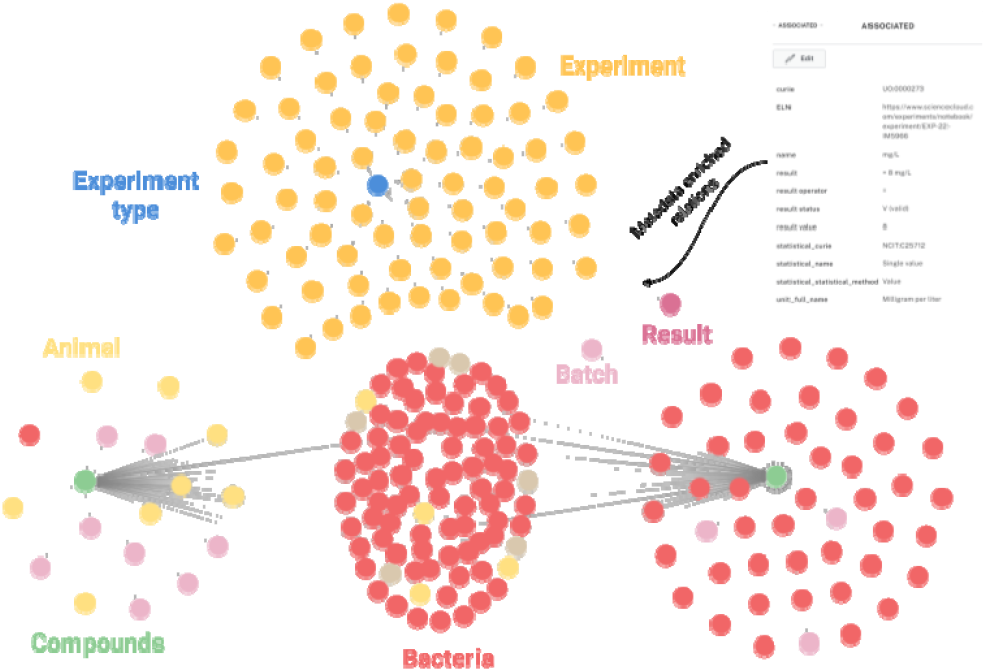
Knowledge graph framework for representation of data of GNA NOW. Here is a screenshot of the Neo4J graph generated through the template showing connectivity between *in-vivo* and *in-vitro* tested compounds.

The FAIRification framework process, as implemented in FAIRplus, is typically planned and implemented over a period of three months. This timeline allows for the detailed breakdown of tasks and helps manage expectations effectively. From the six tasks outlined at the beginning of the Design and Implementation phase of the FAIRification framework **(Figure 2, Step 5)**, we have successfully completed four tasks: a) identification and mapping of the data dictionary with controlled vocabularies and ontologies, b) ensuring the Lab Data Template adheres to tidy principles, c) development of a KG as a viable product for breaking data silos, and d) deposition of the generic KG workflow in a public repository for reuse. However, two remaining tasks— creating a recipe in the FAIR Cookbook and disseminating the GNA NOW data in public repositories—were at risk and could not be completed within the given time frame. Despite these challenges, the progress made on the first four tasks significantly advances our FAIRification goals, laying a solid foundation for future work, where the remaining tasks can be completed. These tasks are currently being dealt with and align together with the project’s timeline to generate impact on data collected and generated in GNA NOW.

### Post-FAIRification assessment

Conducting this analysis offers multiple benefits: it provides a sense of achievement for the efforts dedicated to improvement, keeps the focus on the path of FAIR data management rather than on isolated improvement tasks with no long-term benefit, and demonstrates that FAIR is a continuous journey with tangible steps taken in this direction. Reflecting on our FAIRification improvement through the FAIR-DSM, we found that by implementing the tasks outlined in the previous section, we were able to elevate our maturity from Level 0 to Level 3, as depicted in **Figure 5**. This accomplishment underscores our commitment to advancing data management practices and sets a clear path for future enhancements. The definition of a data dictionary with controlled vocabularies and ontologies as part of the Lab Data Templates served as the main foundation for improvement in the “Content” dimension of our dataset. Additionally, the presence of a structured representation of data and its ability to capture metadata allowed for the improvement in the “Representation” dimension. The development of a modified version of a triple store (i.e. the KG) to connect and host the data distributed across resources provided a platform for efficient retrieval of data and metadata, alongside the unique indexing of each experiment. Furthermore, the inherent capability of database systems such as Neo4J, PostgreSQL, or MySQL to offer programmatic access indirectly elevated the FAIRness level by making the data accessible through multiple channels. Achieving maturity Level 3 in the FAIR-DSM signifies that the datasets are now available in a structured and formalized format. This structured approach enables the datasets to be deposited in public repositories, facilitating reuse by researchers outside the project if needed.

**Figure 5:**
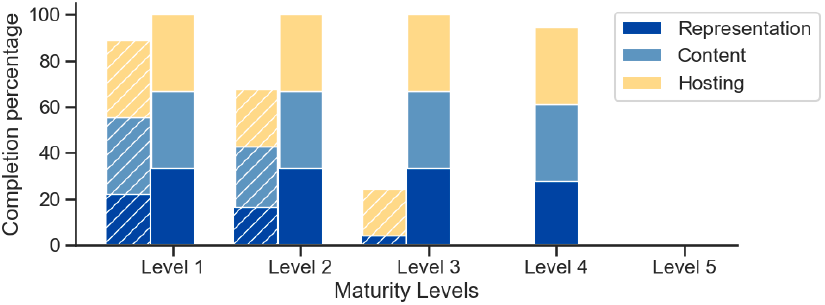
Summary of the pre- and post-FAIRification Assessment of Lab Data Template. The pre-FAIRification assessment results are depicted with dashed bars, while the post-FAIRification assessment results are shown with normal bars. Additionally, the percentage of compliance across the three dimensions of the FAIR-DSM is demonstrated for each level.

## Discussion

FAIR data management is an essential framework for enhancing the utility and efficiency of data in scientific research, especially in drug discovery and development ^36^. In this work, we leveraged the Lab Data Template-oriented recording of experiments to address their machine readability and interoperability alongside electronic lab notebook entries. Templated inputs enable researchers to easily record their data in a metadata-enriched manner without often knowing this is happening ‘under the hood.’ In addition to this, integration across our disparate resources (i.e compound management systems with Grit42 and electronic lab notebooks with Biovia) was accomplished through the generation of data dictionaries. Although widely used in the field, we found that such dictionaries are often insufficiently linked to existing ontologies and controlled vocabularies. We have addressed this gap by ensuring robust connections to relevant ontologies and controlled vocabularies. Another commonly overlooked hindrance to data reuse is the provision of clear usage licenses and detailed provenance information, as shown in this work. Tackling these two issues can facilitate data access and reuse for project collaborators and beyond. Furthermore, we established an interlink between the experimental protocols and experimental results, allowing for tracking of experimental records across silos.

In the GNA NOW project, a group of stakeholders played a crucial role in data management, much like in any other antimicrobial drug discovery project. These stakeholders include researchers who generate data through experiments, data managers who ensure the cataloging of the generated data and their processes, and data scientists who utilize the generated data to make informed decisions for future research. Together, these stakeholders are responsible for the FAIR aspects of the data and are, therefore, the best set of individuals to contribute towards the FAIR data-driven discussion and implementations. These stakeholders were integral to the FAIRification framework used to enhance the FAIRness level of the datasets. Notably, the FAIRification framework is peculiar as it is the only tool that has been developed in this context. Moreover, it has been optimized and improved over time-based on interactions with around 20 IMI projects and is currently being developed and maintained by the ELIXIR Interoperability platform (https://elixir-europe.org/platforms/interoperability). Furthermore, the framework acknowledges the diverse stakeholders involved in the FAIR journey, thereby aiding in organizing the work and identifying what is needed at each stage. Ultimately, the efforts of data FAIRification aim to foster a cultural shift towards better data management practices. This shift will ensure increased data quality and longevity, promoting a more sustainable and effective approach to handling research data.

The FAIRification effort has had a significant impact on the GNA NOW project, leading to numerous achievements that might not have been possible previously. This impact extended to multiple stakeholder groups and was not confined to the data management team. For instance, project and data managers benefited from the KG platform, which served as a “one-stop-shop” resource for summarizing all the experimental efforts within the consortium. The KG provided a holistic view of information and captured relationships between compounds and bacterial behaviour that might have otherwise gone unnoticed. This platform also facilitated efficient scientific reporting, allowing for the analysis of general trends within a single discipline (such as *in-vitro*) and enabling a focused examination of specific datasets across multiple disciplines (*in-vitro* and *in-vivo*) with assurance that all relevant data was included. For medicinal chemists, the ability to query compound scaffolds and review their *in-vitro* and *in-vivo* experimental endpoints collectively enabled the selection and nomination of promising scaffolds for future experiments. Overall, the KG enhanced the visualization and traversal of data distributed across the different infrastructure silos, significantly improving the efficiency and effectiveness of the project. The implementation of Lab Data Templates in GNA NOW not only streamlined data collection but also facilitated their adoption in other IMI/IHI projects within the AMR Accelerator (https://amr-accelerator.eu/), particularly in COMBINE (https://amr-accelerator.eu/project/combine/). Alongside the development of the KG, identifying a suitable private or public repository (such as grit in this case), allowed for the formulation of long-term sustainability strategies for the data beyond the lifetime of the project. In summary, the FAIRification process had a profound impact on the project’s efficiency and effectiveness, benefiting various roles within the consortium and setting a precedent for future data management practices in similar projects.

A major concern in the AMR domain is the reusability of data collected across different dimensions, ranging from surveillance to compound bioactivity. Despite extensive research efforts to combat antimicrobial resistance, the lack of standardization and associated metadata in these datasets hinders their scalable use, particularly through advanced methods like machine learning. One way to address this challenge is by increasing awareness of FAIR principles within the AMR community and showcasing real-world examples of their application and impact. By transitioning towards FAIRer data solutions, researchers can enhance data integration, analysis, and reuse, ultimately accelerating progress in the fight against AMR. The FAIR tools and processes outlined in this work are flexible and thus be applied across various AMR projects. We aim to guide AMR researchers on the practical implementation of FAIR principles using exemplar data from GNA-NOW.

The decision to work with Excel-based Lab Data Templates might seem trivial, but it was the best way to onboard lab scientists and ensure their involvement in the data management process. This approach acknowledged the working practices of lab scientists and resulted in minimal disruption to their daily work or additional training requirements. Moreover, the Lab Data Templates we designed enabled masking of the technical details of FAIR, such as ontology-compliant data dictionaries, thus maintaining its utility without interfering with daily tasks. During our journey, we learned that collaboration between lab scientists, data managers, and data scientists is key to gathering detailed information about experimental lab work and implementing tailored FAIR data management and analysis. Therefore, from our experience, we recommend FAIR efforts that integrate with existing data recording and analysis routines of the lab scientists by utilizing known software tools coupled with a FAIR-compliant workflow from data managers and data scientists. This approach ensures the smooth and uninterrupted flow of data across platforms and machines. In the case of GNA NOW, this was accomplished through the establishment of Lab Data Templates, ontology and controlled vocabulary-based data dictionaries, and the generation of a KG-based platform to understand the connectivity of data distributed across multiple resources. In conclusion, implementing FAIR principles can be challenging due to the need for comprehensive data documentation, infrastructure development, and adherence to community standards. However, the benefits of FAIR data management—enhanced collaboration, improved research efficiency, and greater transparency—are substantial, driving the push for its widespread adoption in the scientific community and beyond.

## Methods

### Lab Data

The lab experiments were performed by North Bristol, NHS Trust, interested readers can obtain the details on standard operating procedures from Marie Attwood. The in vitro microbiology methods (MICs) were based on the latest ISO standards (ISO 20776-1; https://www.iso.org/ics/07.100/x/) where applicable across all sites. Where ISO standards are not stated, EUCAST guidelines (https://www.eucast.org/eucastguidancedocuments) were employed in order to generate robust, reproducible, and high-quality data. Experiments with *in vitro* pH manipulations, altered media compositions, synergy checkerboard assays and time-kill curves methods adhered as closely as possible with EUCAST guidance documents/standards available only with essential minimal deviation in order to maintain reproducibility. Due to the high level of throughput required for this project and the corresponding timeframes, replicates were performed on 10% of all experiments. All methods were rigorously validated prior to experimentation. Bacterial strains were obtained from the clinical collection at North Bristol NHS Trust within the last 3 years. Media used for *in vitro* purposes was Mueller Hinton broth II from BD, which was standardized across multiple sites. *In vivo* experiments were performed in state-registered facilities with ethical clearance. All data components were validated across both *in vitro* and *in vivo* experimentations.

### FAIRification framework and assessment

The FAIRification framework was implemented as described by Welter D. *et al*. ^29^. The templates for the FAIRification framework were taken from Zenodo at https://zenodo.org/records/7501964. The FAIR-DSM assessment tool was hosted online at https://fairdsm.biospeak.solutions/. This tool was used to perform the FAIR assessments for the datasets, and the final report was printed.

### Lab Data templates as a source for ingesting data into grit

The Lab Data Templates, based on Excel spreadsheets, were established to collect *in vitro* and *in vivo* results in a standardized format from different partner sites of the GNA NOW consortium. The goal was to import the content of these completed Lab Data Templates to the projects’ data repository, grit. Excel was chosen because it remains a standard software tool in many research laboratories for the initial recording of experimental parameters and results. The standard format for the Lab Data Template was identified through a survey conducted with the partner sites in GNA NOW. This survey aimed to gain insight into how experiments were conducted and the types of data and results collected. This understanding helped in creating a template that met the specific needs and practices of the researchers involved. Based on the feedback, we first developed a Lab Data Template for the set of *in vitro* experiments. Subsequently, we created an extended version of the template for *in vivo* studies in mice. This approach ensured that the templates were tailored to the specific needs of the researchers and the types of experiments they were conducting. **Table 2** outlines the columns of the Lab Data Template that the researchers need to fill for each experimental study. To ensure consistency in data entry across all laboratories, both Lab Data Templates were linked with data dictionaries. These dictionaries, managed by the project data manager, are separate Excel files containing standardized experimental terms, bacterial strain identifiers, and compound information based on public ontologies (terms) and agreed naming conventions (strains and compounds). Overall, this system helped maintain uniformity and accuracy in the data collected from various sites.

**Table 2:** Data capture in the *in vitro* and *in vivo* Lab Data Template. The sections in the Lab Data Template are divided into four broad groups: a) “Study Detail” capturing the generic experimental study information, b) “Treatment” capturing the compound, bacterial, and animal metadata on which the experiment will be conducted, c) “Activities”, specific to in vivo only, captures the timepoint details for experiments planned, and d) “Experiment Results” capturing the final endpoint results of the single-point (*in-vitro*) and multi-point (*in-vivo*) experiments.

| Group | *In vitro* lab data template | *In vivo* lab data template |
| --- | --- | --- |
| Study Details | STUDYID<br>EXPID<br>SITE<br>PROVENANCE | STUDYID<br>EXPID<br>SITE<br>PROVENANCE<br>STUDY_TYPE<br>PROJECT_LICENCE_NUMBER<br>STUDY_START_DATE<br>STUDY_PROTOCOL_NAME<br>SPECIES_NAME<br>ANIMAL_STRAIN<br>ANIMAL_SEX<br>ANIMAL_AGE_RANGE<br>ANIMAL_BODYWEIGHT_MEAN<br>ANIMAL_BODYWEIGHT_RANGE<br>ANIMAL_VENDOR<br>HOUSING_CAGE_NO_ANIMALS<br>HOUSING_CAGE_SIZE<br>HOUSING_FOOD<br>HOUSING_FOOD_RESTRICTED<br>HOUSING_FOOD_SUPPLEMENT<br>HOUSING_LIGHT_DARK_CYCLE |
| Treatment | CPD_ID<br>EXT_CPD_ID<br>BATCH_ID<br>EXT_BATCH_ID<br>BACTERIAL_STRAIN_NAME<br>BACTERIAL_STRAIN_SITE_REF<br>SPECIES_NAME | CPD_ID<br>EXT_CPD_ID<br>BATCH_ID<br>EXT_BATCH_ID<br>BACTERIAL_STRAIN_NAME<br>BACTERIAL_STRAIN_DOSE<br>GROUP_DESCRIPTION<br>ANIMAL<br>INFECTION_ROUTE<br>PRETREATMENT_CPD_ID<br>PRETREATMENT_BATCH_ID<br>PRETREATMENT_DOSING_INFO<br>PRETREATMENT_DOSE<br>PRETREATMENT_ROUTE_OF_ADMINISTRATION<br>DOSE<br>DOSING_INFO<br>FREQUENCY<br>TDD<br>ROUTE_OF_ADMINISTRATION<br>ANALGESIA_CPD_ID<br>ANALGESIA_BATCH_ID<br>ANALGESIA_DOSING_INFO<br>ANALGESIA_ROUTE_OF_ADMINISTRATION<br>COMMENT |
| Activities | - | GROUP_DESCRIPTION<br>ACTIVITY<br>PLANNED_RELATIVE_TIMEPOINT<br>COMMENT |
| Experiment Results | EXPID<br>EXPERIMENT_DATE<br>EXPERIMENT_TYPE<br>BIOMATERIAL<br>PROTOCOL_NAME<br>RESULT_TYPE<br>STATISTICAL_METHOD<br>RESULT_OPERATOR<br>RESULT_VALUE<br>RESULT_UNIT<br>CONTROL_GROUP<br>RESULT_STATUS<br>COMMENTS<br>NO_OF_REPLICATES<br>MEDIUM<br>MEDIUM_PH<br>MEDIUM_ADDITIVES | EXPID<br>EXPERIMENT_DATE<br>EXPERIMENT_TYPE<br>BIOMATERIAL<br>PROTOCOL_NAME<br>RESULT_TYPE<br>STATISTICAL_METHOD<br>RESULT_OPERATOR<br>RESULT_VALUE<br>RESULT_UNIT<br>CONTROL_GROUP<br>RESULT_STATUS<br>COMMENTS<br>PLANNED_RELATIVE_TIMEPOINT<br>RELATIVE_TIMEPOINT |

### Mapping terms to domain-relevant ontologies and terminologies

We ensured that most of the terms available in the Lab Data Template are standardized, aligned, and integrated with those present in biomedical ontologies. To do so, we used the Ontology Lookup Service (OLS; https://www.ebi.ac.uk/ols4), a data repository of more than 200 biomedical ontologies, to manually identify ontologies for our use case. In OLS, we identified twelve ontologies that could be reused: NCBI Taxonomy (NCBITaxon), Experimental Factor Ontology (EFO), Ontology for Biomedical Investigations (OBI), STATistics Ontology (STATO), NCI Thesaurus (NCIT), BioAssay Ontology (BAO), Uber-anatomy ontology (UBERON), Ontology of Prokaryotic Phenotypic and Metabolic Characters (MicrO), Antibiotic Resistance Ontology (ARO), Units of Measurement Ontology (UO), Phenotype and Trait Ontology (PATO), and Ontology of Biological and Clinical Statistics (OBCS).

Additionally, experimental data requires the preparation of a medium for growth for cultures. Previously, reports for discrepancies in the reproducibility of results due to vendor issues have been raised ^37,38^. To avoid such discrepancies, purchases made by vendor-dedicated sites such as growth mediums were mapped to their respective vendor catalog identifiers, for instance, SIGMA (https://www.sigmaaldrich.com/), for growth medium and ATCC (https://www.atcc.org/), for non-clinical bacterial cultures.

Moreover, the experimentalists commonly used a large number of assay-specific terms in the project. To capture such terminology, we built a project-specific terminology dictionary called GNA NOW. Together, the collection of biomedical ontologies on OLS, vendor catalog identifiers, and project-specific terms formulated the basis for the ontology metadata dictionary for the Lab Data Template. **Table 3** enlists details on the above-mentioned ontologies and their corresponding utility in the Lab Data Template.

**Table 3:** List of ontologies used by the data dictionary of the Lab Data Template.

| Ontology name | Ontology Webpage | Ontology description | Lab Data Template column name |
| --- | --- | --- | --- |
| NCBITaxon | https://www.ncbi.nlm.nih.gov/taxonomy | Ontology for classification and nomenclature of organisms | BIOMATERIAL, SPECIES_NAME |
| EFO | https://www.ebi.ac.uk/efo/ | Ontology for description of experimental variables | EXPERIMENT_TYPE |
| OBI | https://obi-ontology.org/ | Ontology for communicating scientific investigations and results | PRETREATMENT_ROUTE_OF_ADMINISTRATION, ROUTE_OF_ADMINISTRATION, INFECTION_ROUTE |
| STATO | https://stato-ontology.org/ | Ontology for statistical methods and experiments. | STATISTICAL_METHOD |
| NCIT | https://ncit.nci.nih.gov/ncitbrowser/ | Terminology catalog for clinical care, translational research, and administrative activities. | BIOMATERIAL, EXPERIMENT_TYPE, MEDIUM, STATISTICAL_METHOD |
| BAO | http://bioassayontology.org/ | Ontology describing chemical biology screening assays and their results | EXPERIMENT_TYPE |
| UBERON | https://obophenotype.github.io/uberon/ | Ontology for cross-species anatomy | MEDIUM |
| MicrO | https://github.com/carrineblank/MicrO | Ontology of microbiological terms | MEDIUM |
| ARO | https://card.mcmaster.ca/browse | Ontology of known antibacterial resistance determinants and associated antibiotics | EXPERIMENT_TYPE, MEDIUM |
| UO | https://github.com/bio-ontology-research-group/unit-ontology | Ontology of units of measurements | RESULT_UNIT |
| PATO | https://pato-ontology.github.io/pato/ | Ontology of phenotypes | ANIMAL_SEX |
| OBCS | https://github.com/obcs/obcs | Ontology covering biological and clinical statistics | STATISTICAL_METHOD |
| SIGMA | https://www.sigmaaldrich.com/ | Assay material commercial vendor | MEDIUM |
| ATCC | https://www.atcc.org/ | Microorganisms, cell lines, and other materials commercial vendor | BACTERIAL_STRAIN_NAME, ANIMAL_STRAIN |
| GNA NOW | https://github.com/IMI-COMBINE/GNA-NOW/blob/main/data/mapping_files/gna_ontology.tsv | Project-specific ontology that contains terms not found in any public ontology | EXPERIMENT_TYPE, ANIMAL_VENDOR, RESULT_UNIT, STUDY_TYPE |

### Knowledge graph construction

We developed our graph database through a meticulously planned process, which involved the integration of data derived from both *in-vivo* and *in-vitro* experiments. Opting for the robust infrastructure offered by Neo4J (https://neo4j.com/), we transformed our data into property graphs within this environment. Leveraging the capabilities of the Python plugin py2neo (https://pypi.org/project/py2neo/), we efficiently constructed and executed queries on the underlying property graph within Neo4J. Additionally, recognizing the diverse skill sets within our consortium, we used the Neo4J Bloom instance to facilitate text-based searches for those less familiar with graph databases. This user-friendly interface empowered all members to access and explore the data, irrespective of their expertise in graph theory or database management.

## Data availability

The Data Templates, along with the metadata dictionaries using consistent terminologies and ontologies, are available on Zenodo^39^ for reuse. The datasets that were mapped to this data template as a use case will not be made public since it is a proprietary dataset. However, we have provided a dummy template for both experiments to allow users to reproduce the workflow independently.

## Code availability

The source code for processing the template files and converting them into knowledge graphs has been made available on GitHub at https://github.com/IMI-COMBINE/template2graphs. This repository includes documentation for reusing functions and workflows for building the KG, provided that the underlying data is compliant with the Lab Data Template. Due to data restrictions, the data and the graph instance will not be made public.

## Acknowledgments

We would like to thank Andrea Zaliani for reviewing the manuscript and providing us with valuable feedback. This work and the authors were primarily funded by the following projects: FAIRplus (IMI 802750), COMBINE (IMI 853967), and GNA NOW (IMI 853979).

## Author contribution

Y.G: Conceptualization, Supervision, Methodology, Writing - Original draft preparation and Visualization; T.A.D: Conceptualization, Methodology, Writing - Original draft preparation and Visualization; G.W: Supervision, Methodology and Writing - Original draft preparation and Visualization; V.I: Conceptualization and Writing - Original draft preparation and Visualization; C.S.K: Writing - Reviewing and Editing; N.J: Writing - Reviewing and Editing; M.K: Writing - Reviewing and Editing; M.A: Writing - Reviewing and Editing; P.G: Writing - Reviewing and Editing.

## Competing interests

None

## Notes

https://doi.org/10.5281/zenodo.12720580

https://github.com/IMI-COMBINE/template2graphs

### Competing Interest Statement

The authors have declared no competing interest.

### Summary of Updates

Version has been updated based on comments from the reviewers.

## References

1. Jacobsen, A. et al. FAIR Principles: Interpretations and implementation Considerations. Data Intelligence 2, 10–29 (2020).

2. Publications Office of the European Union. European Research Data Landscape□: final report. Publications Office of the EU https://op.europa.eu/en/publication-detail/-/publication/03b5562d-6a35-11ed-b14f-01aa75ed71a1 (2022).

3. Kruhse-Lehtonen, U. & Hofmann, D. How to define and Execute your data and AI Strategy. Harvard Data Science Review (2020) doi:10.1162/99608f92.a010feeb.

4. Steffens, S. et al. The challenges of research data management in cardiovascular science: a DGK and DZHK position paper—executive summary. Clinical Research in Cardiology 113, 672–679 (2023).

5. Murphy, F. Open access, open data, FAIR Data and their implications for life sciences researchers. Emerging Topics in Life Sciences 2, 759–762 (2018).

6. Coles, S. J., Frey, J. G., Willighagen, E. & Chalk, S. Taking FAIR on the CHIN: the Chemistry Implementation Network. Data Intelligence 2, 131–138 (2020).

7. Iturbide, M. et al. Implementation of FAIR principles in the IPCC: the WGI AR6 Atlas repository. Scientific Data 9, (2022).

8. Wise, J. et al. Implementation and relevance of FAIR data principles in biopharmaceutical R&D. Drug Discovery Today 24, 933–938 (2019).

9. Dumit, V. I. et al. From principles to reality. FAIR implementation in the nanosafety community. Nano Today 51, 101923 (2023).

10. Vesteghem, C. et al. Implementing the FAIR Data Principles in precision oncology: review of supporting initiatives. Briefings in Bioinformatics 21, 936–945 (2019).

11. Forschungsgemeinschaft, D. Guidelines for safeguarding good research practice. Code of conduct. Zenodo (CERN European Organization for Nuclear Research) (2022) doi:10.5281/zenodo.6472827.

12. Bloemers, M. & Montesanti, A. The FAIR Funding Model: Providing a Framework for Research Funders to Drive the Transition toward FAIR Data Management and Stewardship Practices. Data Intelligence 2, 171–180 (2020).

13. European Commission, Directorate-General for Research and Innovation [Maxwell, L]. Maximising Investments in Health Research: FAIR Data for a Coordinated COVID-19 Response□: Workshop Report. (Publications Office of the European Union, 2022). doi:10.2777/726950.

14. David, R. et al. “Be sustainable”: EOSC□Life recommendations for implementation of FAIR principles in life science data handling. EMBO Journal 42, (2023).

15. Gadiya, Y. et al. FAIR data management: what does it mean for drug discovery? Frontiers in Drug Discovery 3, (2023).

16. Alharbi, E. et al. Selection of data sets for FAIRification in drug discovery and development: Which, why, and how? Drug Discovery Today 27, 2080–2085 (2022).

17. Van Vlijmen, H. et al. The need of industry to go FAIR. Data Intelligence 2, 276–284 (2020).

18. Alharbi, E., Skeva, R., Juty, N., Jay, C. & Goble, C. A FAIR-Decide framework for pharmaceutical R&D: FAIR data cost–benefit assessment. Drug Discovery Today 28, 103510 (2023).

19. Wise, J. et al. Implementation and relevance of FAIR data principles in biopharmaceutical R&D. Drug Discovery Today 24, 933–938 (2019).

20. Alharbi, E., Skeva, R., Juty, N., Jay, C. & Goble, C. Exploring the current practices, costs and benefits of FAIR implementation in pharmaceutical research and Development: a qualitative interview study. Data Intelligence 3, 507–527 (2021).

21. Vesteghem, C. et al. Implementing the FAIR Data Principles in precision oncology: review of supporting initiatives. Briefings in Bioinformatics 21, 936–945 (2019).

22. Harrow, I., Balakrishnan, R., McGinty, H. K., Plasterer, T. & Romacker, M. Maximizing data value for biopharma through FAIR and quality implementation: FAIR plus Q. Drug Discovery Today 27, 1441–1447 (2022).

23. Shabani, M. & Obasa, M. Transparency and objectivity in governance of clinical trials data sharing: Current practices and approaches. Clinical Trials 16, 547–551 (2019).

24. Ghiandoni, G. M., Evertsson, E., Riley, D., Tyrchan, C. & Rathi, P. C. Augmenting DMTA using predictive AI modelling at AstraZeneca. Drug Discovery Today 103945 (2024) doi:10.1016/j.drudis.2024.103945.

25. Brochu, F. et al. Federation of Imaging Data for Life Sciences Current Status of Metadata Collection for High Content Screening, Mass Spectrometry Imaging and Light Sheet Microscopy of AstraZeneca, GlaxoSmithKline and NPL. http://eprintspublications.npl.co.uk/8838/1/MS24.pdf (2020) xdoi:10.47120/npl.ms24.

26. Kelm, J. M., Ferrer, M., Bittner, M.-I. & Lal□Nag, M. Data standards in drug discovery: A long way to go. Drug Discovery Today 29, 103879 (2024).

27. Rocca-Serra, P. et al. The FAIR Cookbook - the essential resource for and by FAIR doers. Scientific Data 10, (2023).

28. Beyan, O. et al. D2.5 FAIRPlus FAIR Data Maturity Framework. Zenodo (CERN European Organization for Nuclear Research) https://zenodo.org/records/5040592 (2021) xdoi:10.5281/zenodo.5040592.

29. Welter, D. et al. FAIR in action - a flexible framework to guide FAIRification. Scientific Data 10, (2023).

30. Da Silva Santos, L. O. B., Burger, K., Kaliyaperumal, R. & Wilkinson, M. D. FAIR Data Point: A FAIR-Oriented approach for metadata publication. Data Intelligence 5, 163–183 (2023).

31. Corcho, O. et al. EOSC Interoperability Framework□: Report from the EOSC Executive Board Working Groups FAIR and Architecture. (2021). doi:10.2777/620649.

32. Wilkinson, M. D. et al. FAIR Assessment Tools: Towards an “Apples to Apples” Comparisons. Zenodo (CERN European Organization for Nuclear Research) https://zenodo.org/records/7463421 (2022) xdoi:10.5281/zenodo.7463421.

33. Cooper, M. A. A community-based approach to new antibiotic discovery. Nature Reviews. Drug Discovery 14, 587–588 (2015).

34. Reimer, L. C. et al. BacDive in 2022: the knowledge base for standardized bacterial and archaeal data. Nucleic Acids Research 50, D741–D746 (2021).

35. Wickham, H. Tidy data. Journal of Statistical Software 59, (2014).

36. Ugochukwu, A. I. & Phillips, P. W. B. Open Data Ownership and Sharing: Challenges and opportunities for application of FAIR principles and a Checklist for data managers. Journal of Agriculture and Food Research 16, 101157 (2024).

37. Williams, M. Reagent validation to facilitate experimental reproducibility. Current Protocols in Pharmacology 81, (2018).

38. Jaric, I. et al. Using mice from different breeding sites fails to improve replicability of results from single-laboratory studies. Lab Animal 53, 18–22 (2023).

39. Witt, G. et al. Supplementary data files for manuscript titled “From spreadsheet lab data templates to knowledge graphs: A FAIR data journey in the domain of AMR research”. Zenodo (CERN European Organization for Nuclear Research) https://zenodo.org/records/12720580 (2024) xdoi:10.5281/zenodo.12720579.

